# Three-dimensional regulation of *HOXA* cluster genes by a *cis*-element in hematopoietic stem cell and leukemia

**DOI:** 10.1101/2020.04.16.017533

**Authors:** Xue Qing David Wang, Haley Gore, Pamela Himadewi, Fan Feng, Lu Yang, Wanding Zhou, Yushuai Liu, Xinyu Wang, Chun-wei Chen, Jianzhong Su, Jie Liu, Gerd Pfeifer, Xiaotian Zhang

## Abstract

Proper gene regulation is crucial for cellular differentiation, and dysregulation of key genes can lead to diseased states such as cancer. The HOX transcription factors play such a role during hematopoiesis, and aberrant expression of certain *HOXA* genes is found in certain acute myeloid leukemias (AMLs). While studies have shown that these genes are targeted by a variety of mutant proteins including mutant NPM1, MLL fusions, and NUP98 fusions, little is known about how long-range 3D chromatin interactions regulate the *HOXA* genes in normal hematopoiesis and leukemia. Here, we report the interaction between the *HOXA* cluster with a ∼1.3 Mb upstream DNA methylation Canyon termed “*Geneless Canyon*” (*GLC*) in human CD34+/CD38-hematopoietic stem cells (HSCs) and AML cell lines. We show that CRISPR-Cas9 mediated deletion of the whole *GLC* region reduces the expression of the distal *HOXA* genes and compromises HSC and leukemia cells self-renewal. This long-range chromatin interaction brings the *HOXA* cluster in contact with the nuclear pore complex (NPC) at the nuclear periphery, which promotes *HOXA* gene expression and maintains HSC and leukemia cell self-renewal. These findings reveal how long-range 3D chromatin organization regulates key transcription factor genes in both normal and diseased hematopoietic cells.

## Main Text

Hematopoietic stem cells (HSCs) give rise to a wide variety of blood cells through a tightly regulated differentiation process, in which the *HOX* transcription factors play a key role^1,2^. Several *HOXA* genes have been shown to maintain a high level of stem cell self-renewal at the early stages of hematopoiesis, but these genes are repressed as the cells differentiate into specific lineages^3^. Overexpression of these *HOXA* genes is found in several acute myeloid leukemias (AML), where the genes are often directly targeted by chimeric proteins such as MLL-fusions and NUP98-fusion^4,5^. Recent studies in mouse limb development have identified several enhancers interacting with the *HOXA* locus, contributing to their gene regulation during development^6,7^. As of yet, little is known about the role of 3D chromatin organization during hematopoiesis and whether distant enhancers are involved in the regulation of the *HOXA* genes in both normal and diseased states.

The *HOXA* gene cluster is located between two topologically associating domains (TADs) and acts as a boundary for both 5’ and 3’ TADs. At the 5’ end (centromeric), there is a ∼1 Mb TAD encompassing the *HIBADH, TAX1BP1* and *JAZF1* genes, while the 3’ (telomeric) TAD spans ∼1.3 Mb and includes several subTADs formed around genes (Figure 1a). We previously identified an interaction between the *HOXA* locus and an upstream DNA methylation canyon, designated *Geneless Canyon* or *GLC*, in cord blood CD34+ HSPCs^8^. This long-range interaction anchors the 3’ TAD despite a lack of convergent CTCF binding sites (Supplemental Figure S4). Unlike CTCF-mediated loops, the interaction between *GLC* and the *HOXA* locus is broad and covers regions enriched in polycomb-silenced H3K27me3 (Figure 1a, b). The *GLC* also acts as a boundary to an 3’ upstream TAD featuring a gene desert (Figure 1a). We speculated that the *GLC* could function as a *cis*-regulatory element to the posterior *HOXA* genes in addition of being a TAD boundary element. CRISPR-Cas9 mediated deletion of the *GLC* in cord blood CD34+ HSPCs using two flanking guide RNAs yielded a heterogeneous population of *GLC*-deleted and wild-type cells (Figure 1c, d). This population of cells differentiated more rapidly in *ex vivo* culture conditions compared to control cells edited for an off-target deletion (Figure 1e). Furthermore, the *GLC*-deleted cells yielded smaller and significantly lesser number of colonies in a colony formation assay (Figure 1f, g). This suggests that the *GLC* is a non-coding region with functional role in HSPC self-renewal activity. *In situ* Hi-C revealed that partial deletion of the *GLC* resulted in a significant loss of interaction with the *HOXA* locus, particularly at the H3K27me3 enriched regions (Figure 1h). Meanwhile, the active *HOXA* region enriched in H3K27ac (*HOXA6-HOXA10*) also lost interactions with the H3K27ac marked upstream enhancers in *SKAP2* and *SNX10* region^9^ (Figure 1h, i, Supplemental Figure S1b, c). Consistent with the loss of this active enhancer-promoter interaction, the loss of *GLC* also resulted in a significant decreased expression of the active *HOXA7-10* genes and a modest decrease of the interacting *HOXA11* and *A13* (Figure 1j). These results suggest that the *GLC* does not act as a repressor of the distal *HOXA* genes despite its interaction with the H3K27me3 rich region of the locus. Instead, this long-range interaction serves as an activating element for the *HOXA* genes in HSPCs potentially by maintaining promoter-enhancer interaction for the active *HOXA* genes (Figure 1k).

**Figure 1.**
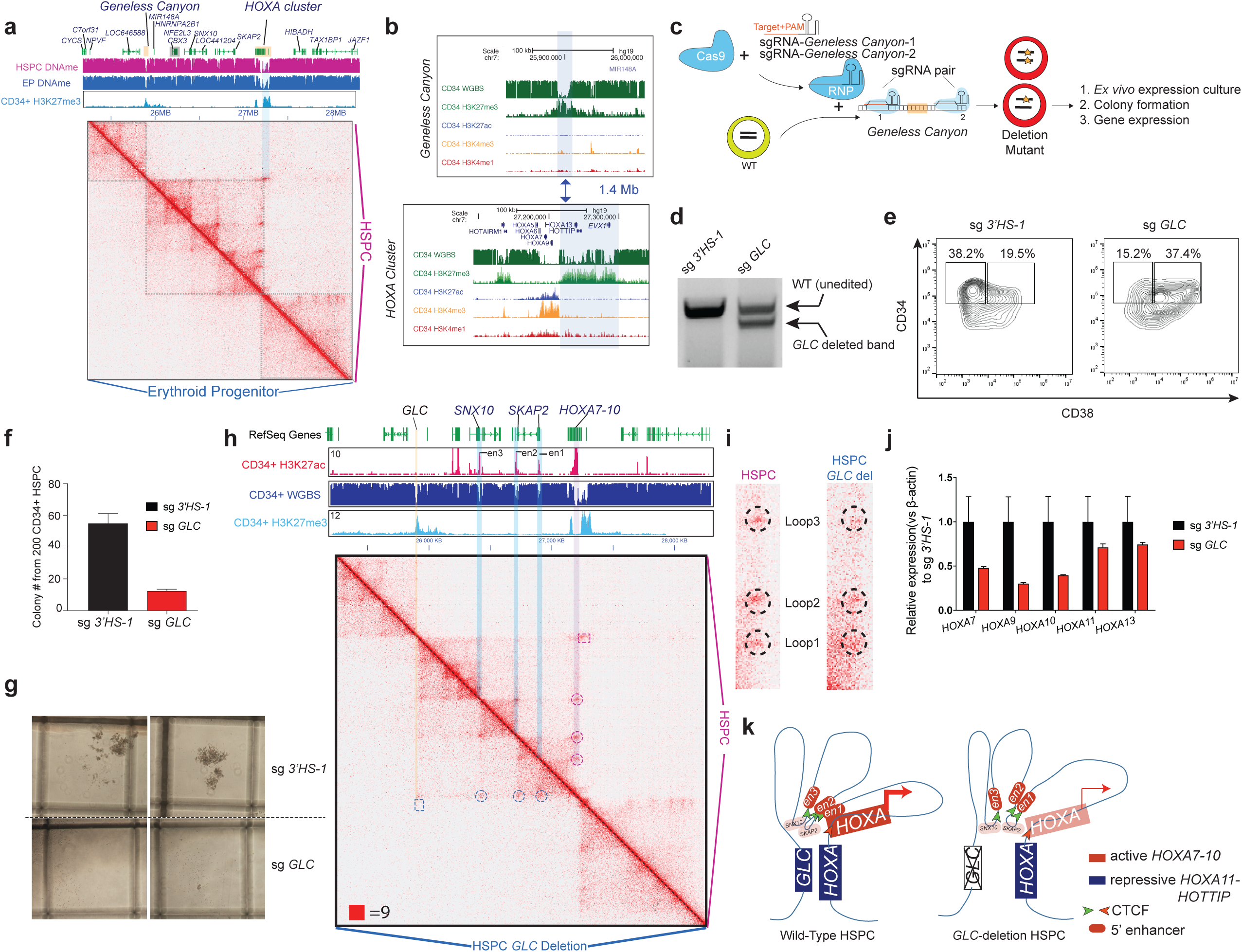
*Geneless Canyon* maintains active stem cell activity and *HOXA* gene expression in human HSPC. **a**). Hi-C contact heatmap of CD34+ HSPC (upper triangle) and differentiated erythroid progenitors (lower triangles). Grey box annotates the 3 TADs around *HOXA* region in both cell types. **b**). The epigenetic state of *HOXA* cluster and *GLC* in CD34+ HSPCs. DNA methylation, H3K27me3, H3K27ac, H3K4me3 and H3K4me1 is shown. **c**). Scheme of using 2 guideRNAs for the deletion of *GLC* in CD34+ HSPCs. **d**). The deletion of *GLC* by paired guideRNA after two rounds of RNP electroporation. **e**). Flow cytometry plot of HSPCs after *GLC* deletion with CD34-FITC and CD38-PE. Flowcytometry is performed after 4 days of *ex vivo* culture in *ex vivo* expansion medium. **f**). Colony number of CD34+ HSPCs electroporated with paired guide RNA targeting hemoglobin locus 3’HS-1 CTCF and *GLC*. n=3, mean ± s.d. is shown. **g**). Colony size of CD34+ HSPCs electroporated with paired guide RNA targeting 3’HS-1 and *GLC*. Grid size is 1cm × 1cm. **h**). Hi-C contact heatmap of CD34+ HSPC and *GLC*-deleted C34+ HSPC (upper/lower triangles, respectively) with overlay of H3K27ac and WGBS. **i**). Zoom-in of *HOXA* interactions with upstream enhancer elements. **j**). *HOXA* gene expression after the deletion of 3’HS-1 CTCF and *GLC*. n=3, mean ± s.d. is shown. **k**). Model of 3D genomic interactions in *HOXA* locus in HSPC.

The *HOXA* transcription factors are often expressed in subtypes of human acute myeloid leukemias (AML) and overexpression of the *HOXA9* gene has been shown to directly transform mouse bone marrow cells^10^. To determine whether the *GLC* plays a role in maintaining the aberrant expression of *HOXA* genes in leukemia, we performed *in situ* Hi-C on primary human leukemia blasts from patients carrying the *MLL-AF9* translocation (AML-4943), the *NPM1* mutation (AML-5577, AML-6527), as well as *HOXA* high expressing cell lines OCI-AML3, MV4-11 and the *HOXA* low expressing cell line Kasumi-1 (Figure 2a, b, Supplemental Figure S5a-d)^11,12^. The AML blasts were mostly CD33+ CD34- (Supplemental Figure S2a-b). The high-resolution Hi-C maps were validated through identification of known genomic translocations such as t(9:11) in AML-4943 blast, t(4:11) in MV4-11 and t(8:21) in Kasumi-1 cells (Supplemental Figure 3a-f). We observed that the genome-wide chromosomal loop interactions were consistent in all 3 AML blasts by aggregate peak analysis (Supplemental Figure S2c)^13^. However, up to 80% of grand canyon interactions were lost in primary AML blasts and almost all such interactions were lost in cell lines (Supplemental Figure S2c, d). In tandem, we performed ChIP-seq on H3K27me3 and H3K27ac histone marks to identify regions of silent and active chromatin in these cells. Overlaying Hi-C contact maps with the histone marks revealed a pattern between the 3D chromatin organization and the underlying epigenome. Strong *GLC* to *HOXA* interaction was observed in the AML-4943 patient blast cells, similar to cord blood CD34+ HSPC with comparable silent chromatin states at both *GLC* and *HOXA* regions (Figure 1a, 2a, c, Supplemental Figure 1d). Additionally, entire *SKAP2* gene is marked by a very broad H3K27ac peak and forms a stripe interaction with the active *HOXA* genes, which is distinct from other cell types (Figure 2a, c). In in AML-5577, the site of contact at the *HOXA* locus shifts patient blasts to the location of matching activating H3K4me3 marks (Supplemental Figure 5a, e). This is also observed in the OCI-AML3 cell line, which is depleted of H3K27me3 at the *GLC*, but strong H3K27ac at both *GLC* and *HOXA* seem to demarcate this long-range interaction (Figure 2b, d). Since both AML-5577 and OCI-AML3 carry the NPM1^W288fs^ mutation, these results suggest that mutant NPM1 may be associated with the structural alteration at the *HOXA* cluster. This long-range interaction was significantly weaker in MV4-11 cells, which showed no H3K27me3 and weak H3K27ac at the *GLC*, and a strong but narrow enrichment of H3K27ac at the active *HOXA*9-A10 genes (Supplemental Figure S5b, e). Meanwhile, the low *HOXA* expressing Kasumi-1 cell line had *HOXA* interaction with convergent CTCF sites downstream of the *GLC* (Figure 2e, Supplemental Figure S5c). This result is particularly interesting as Kasumi-1 cells feature enrichment of H3K27me3 at both *GLC* and distal *HOXA* regions, yet they do not directly form a long-range loop. Taken together, these results show that the *GLC*-*HOXA* interaction occurs primarily in cells expressing high levels of *HOXA* genes, and this interaction is associated by co-occurring histone modifications at both loci (Figure 2f, g).

**Figure 2.**
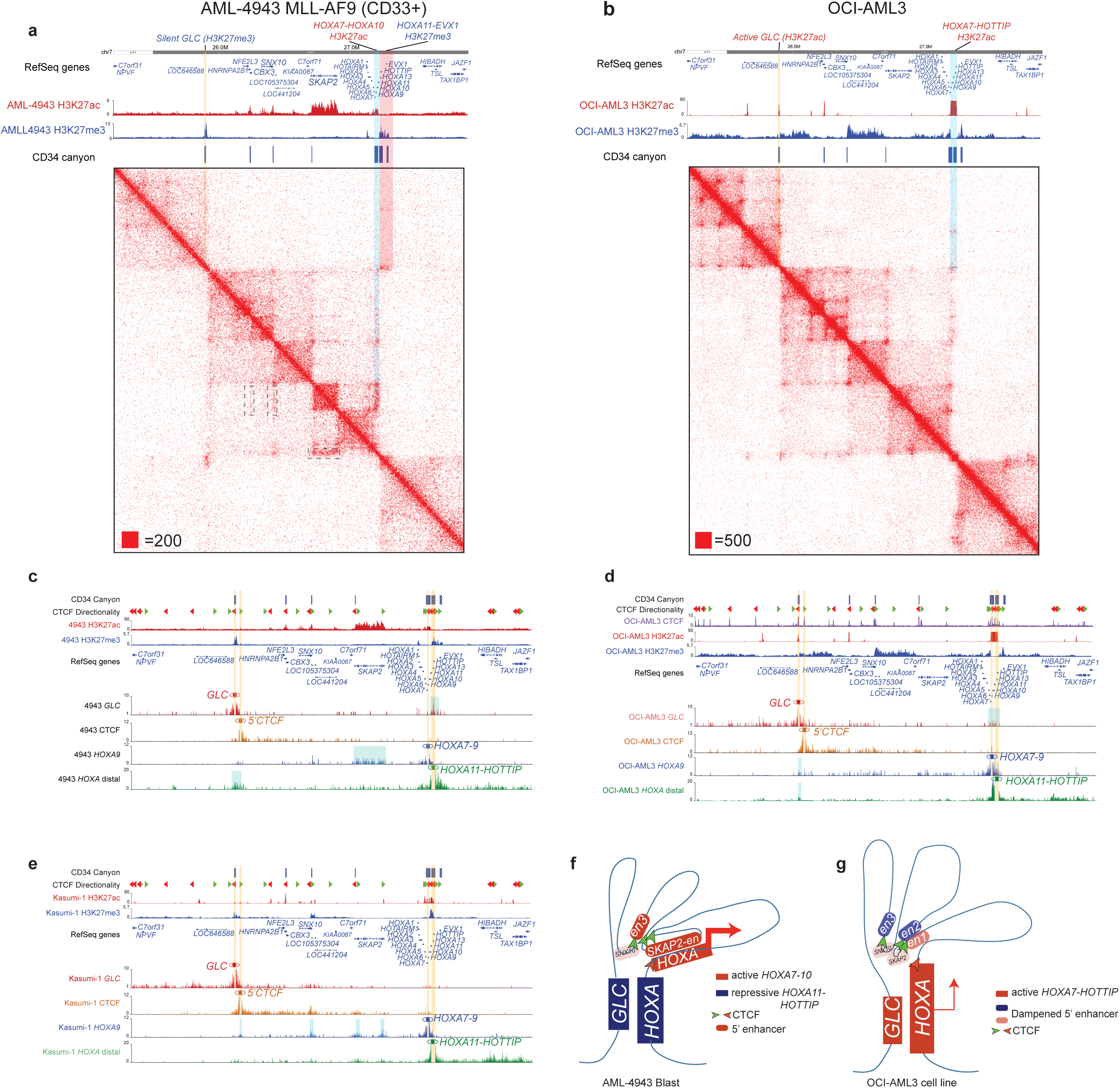
*Geneless Canyon* interacts with HOXA cluster in primary leukemia samples and cell lines. **a**). The contact heatmap in 3 TAD regions in around *HOXA* cluster in primary leukemia blast AML4943. AML4943 carries the *MLL-AF9* translocation. Yellow shade indicates the interacting regions of *GLC* and corresponding *HOXA* regions (*HOXA11* to *EVX1*). Blue shade indicates the active *HOXA* region (*HOXA7-HOXA10*). Grey box annotates the interaction between *SKAP2* and *HOXA7-10* region (horizonal); *SNX10, CBX10* (Vertical). **b**). The contact heatmap in 3 TAD regions in around *HOXA* cluster in OCLI-AML3 cell lines. Yellow shade indicates the interacting regions of *GLC* and corresponding *HOXA* (*HOXA7-HOTTP*) regions. All Hi-C contact map is shown with the corresponding histone ChIP-seq tracks and CD34+ HSPC canyon and whole genome bisulfite sequencing data for the location of *GLC*. **c**). The virtual 4C plot with multiple viewpoints from *GLC*, 5’CTCF, *HOXA7-HOXA10, HOXA11-HOTTIP* in the CD33+ AML-4943 Blast. **d**). The virtual 4C plot with multiple viewpoints from *GLC*, 5’CTCF, *HOXA7-HOXA10, HOXA11-HOTTIP* in the OCI-AML3 cell line. **e**). The virtual 4C plot with multiple viewpoints from *GLC*, 5’CTCF, *HOXA7-HOXA10, HOXA11-HOTTIP* in the Kasumi-1 cell line. Yellow bars highlight the viewpoint regions and blue bars highlight the interacting regions with the specific viewpoint. **f-g**). 3D genomic interaction model of 3’ TAD and HOXA cluster in AML-4943 blast (**f**) and OCI-AML3 cell lines(**g**).

As the *GLC* interacts with active *HOXA* locus in leukemia cells, we used paired sgRNAs to delete the *GLC* in OCI-AML3 and MV4-11 cell lines to determine whether it possesses similar regulatory function as in HSPC (Figure 3a). The CRISPR-Cas9 mediated-deletion created a heterogeneous population of *GLC*-deleted and wild-type cells which were cultured up to 21 days in a growth competition assay. Comparing the two populations at days 7 and 21, the *GLC*-deleted cells declined over time, indicating that the non-coding region is required for the growth of both cell lines (Figure 3b). As the *GLC* spans a ∼17 kb region, we used a CRISPR-Cas9 saturation mutagenesis screen to identify distinct elements within this region that are crucial for cell growth (Figure 3c, Supplemental Figure S6a). We have identified that sgRNA targets within the *GLC* that drop-off over a period of 15 days as well as positive control (Supplemental Figure S6b). The data suggests there are smaller regions in *GLC* required for maintaining cell growth. Comparing the results from both OCI-AML3 and MV4-11 cells, two hot spots were identified at CTCF binding sites within the *GLC* (Figure 3d, e). Curiously, these two CTCF sites are non-convergent with respect to the CTCF sites within the *HOXA* cluster (Supplemental Figure S4). Individual deletion of the CTCF binding sites (designated CTCF-1 and CTCF-2) in cord blood CD34+ HSPC yielded different levels of effectiveness, yet both were found to diminish HSPC self-renewal (Figure 3f, g). Furthermore, several *HOXA* genes and the transcription factor *MEIS1* were downregulated following the loss of CTCF-1 (Figure 3h). Deletion of both CTCF-1 and −2 in OCI-AML3 cell line revealed that loss of either sites could not compete in growth against wild-type cells (Figure 3i). Loss of CTCF-1, being a more efficient deletion, revealed a similar loss of *HOXA* and *MEIS1* expression in this cell line (Figure 3j). These results suggest that, despite the divergent orientation of CTCF binding sites within the *GLC*, these elements have a functional role in regulating the *HOXA* genes. Given the *GLC* switched to active state marked by H3K27ac in the MV4-11 and OCI-AML3 cell lines (Supplemental Figure S6d), the two CTCFs may play a role in maintaining the active expression of *HOXA* genes by stabilizing promoter-enhancer interactions.

**Figure 3.**
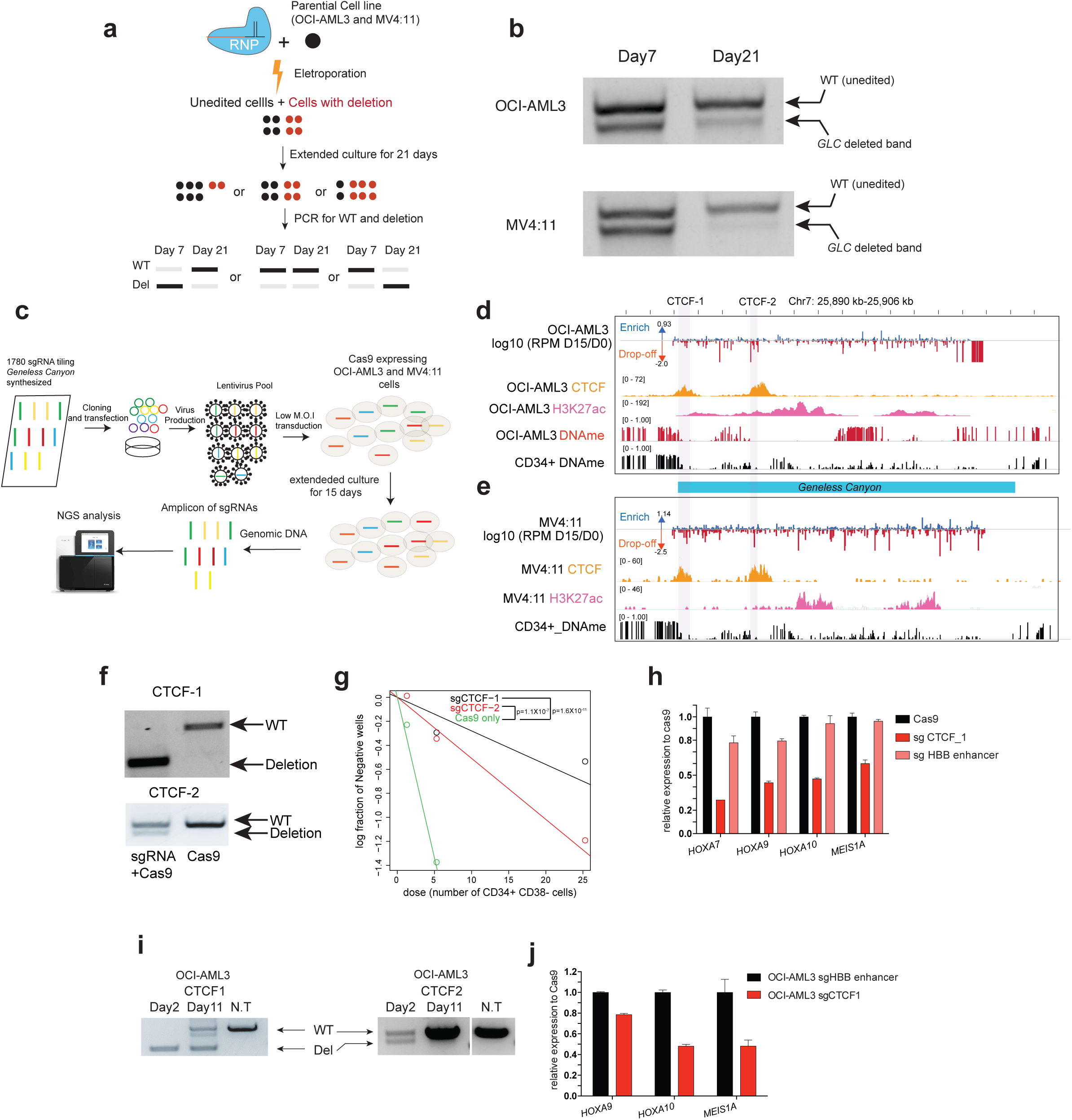
*Geneless Canyon* and CTCF sites within is essential to leukemia cell line maintenance and *HOXA* gene expression. **a**). The scheme of competition assay between *GLC* deletion and WT cell lines with CRISPR/Cas9. **b**). The gel picture of *GLC* deletion and wild type band in cell competition assay for OCI-AML3 and MV4-11 cells. **c**). Scheme for the tilling CRISPR/Cas9 mutagenesis for *GLC.* **d**). The mapping of guide RNA drop-off and enrichment score (calculated as log10(normalized count_day15_/normalized count_day0_)) in OCI-AML cells. OCL-AML3 CTCF, H3K27ac and DNA methylation is shown in parallel **e**). The mapping of guide RNA drop-off and enrichment score (calculated as log10(normalized count_day15_/normalized count_day0_)) in MV4-11 cells. MV4-11 CTCF, H3K27ac is shown in parallel. **f**). Gel picture of CTCF1 and CTCF2 deletion with paired guide RNA in cord blood CD34+ HPSC. **g**). Serial dilution colony forming assay with Cas9, CTCF1-deletion and CTCF2-deletion CD34+ HSPCs. **h**). qPCR detection of active *HOXA* gene expression in CD34+ HSPCs, n=3, mean ± s.d. is shown. **i**). The gel picture of CTCF-1 and CTCF-2 deletion and wild type band in cell competition assay for OCI-AML3 and MV4-11 cells. **h**). qPCR detection of *HOXA9, HOXA10* and *MEIS1A* gene expression in OCI-AML cell line, n=3, mean ± s.d. is shown.

The orientation of CTCF motifs is a key factor in the formation of TADs and chromosomal loops^14^. As both CTCF-1 and −2 are in divergent orientation from the *HOXA* CTCF sites, we investigated whether they formed other TADs elsewhere on the chromosome. We observed a chromosomal loop interaction between CTCF-1 and −2 and upstream (telomeric) CTCF sites (Figure 4a). This spanning region includes a single long non-coding RNA and has been identified as a lamina-associated domain (LAD) in IMR90 cells (Figure 4a)^15^. Additional analysis also shows that the long DNA methylation canyon is likely to be flanked by LAD regions (Supplemental Figure 7a, b). Given the features of constitutive LADs (cLAD), which include low gene density, we surmised that this region would remain as a LAD in HSPC and leukemia cells^16^. Fluorescent *in situ* hybridization (FISH) revealed that both distal *HOXA* and *GLC* were located at the nuclear periphery and remained in proximity to each other in cord blood CD34+ HSPCs (Figure 4b). Furthermore, the *GLC* was found to be close to the network of Lamin B1 located at the inner nuclear membrane in OCI-AML3 cells (Figure 4c). As such, we propose a model in which the *HOXA* locus is spatially tethered to the nuclear periphery through its interaction with the *GLC* and LAD either through repressive or activating domains (Figure 4d, e).

**Figure 4.**
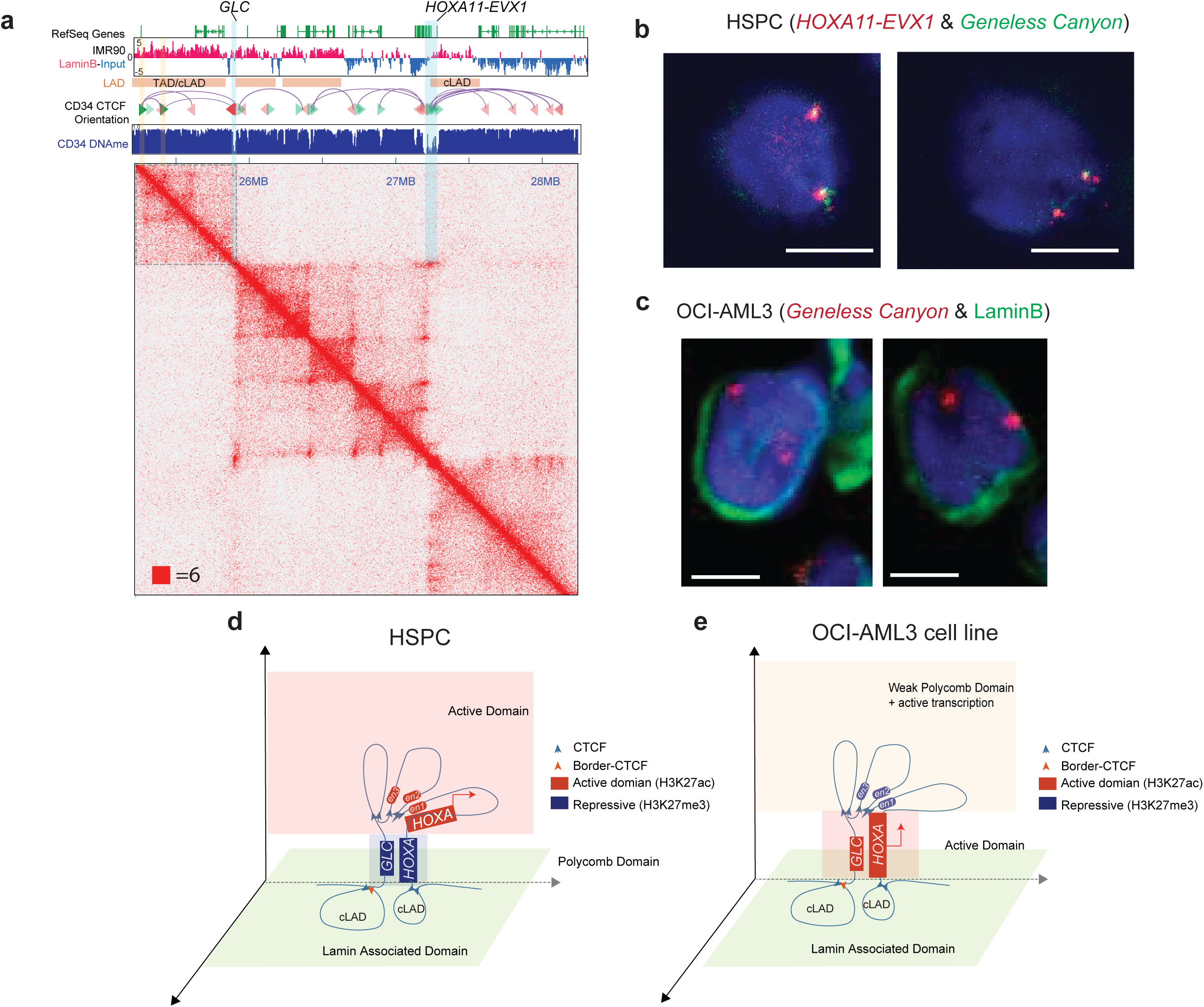
*Geneless Canyon-HOXA* interaction is located at nuclear periphery. **a**). The distribution of Lamin associated domain around *HOXA* cluster. **b**). FISH image of *GLC* (green) and *HOXA* (red) in the cord blood CD34+ cells **c**). Immuno-FISH image of *GLC* (red) and LNMB (green) in OCI-AML3 cells. **d-e**). Model of *HOXA* cluster genomic locus in 3D in normal HSPC (**d**) and OCI-AML3 leukemia cell lines (**e**).

Our data show that the *HOXA* gene cluster interacts with the upstream *Geneless Canyon*, forming a long-range chromosomal loop tethered to the nuclear periphery of HSPCs and cell lines expressing high levels of the *HOXA* transcription factors. While the *GLC* is a polycomb target in CD34+ HSPCs, we observed that it can change epigenetic states across various tissues and cell lines. This is consistent with previous works that show how polycomb targets in early development can act as enhancers in *Drosophila* and mouse models^17-20^. We have previously shown that ablation of the silencing histone marks was sufficient to disrupt the interaction between *GLC* and *HOXA* and alters gene expression at the locus in cord blood CD34+ HSPC (Zhang et al. *Molecular Cell in press*). Here, we observe that the loss of *GLC* significantly hampers the self-renewal capability of CD34+ HSPCs and stunts growth of OCI-AML3 leukemia cell line model. These phenotypes are accompanied by significant downregulation of key *HOXA* genes. Interestingly, the long-range interaction between *GLC* and *HOXA* locus is often mediated at silencing H3K27me3 regions, yet loss of this interaction does not activate the silenced distal *HOXA11* to *HOTTIP* genes. This suggests that the interaction itself may not be altering the epigenetic state at the *HOXA* cluster, but rather play a role in establishing a scaffold in 3D nuclear space that promotes *HOXA* gene expression. This idea has also been echoed by several recent studies with the evidence that Polycomb target loci can act as permissive 3D genomic structure to allow the active transcription of Homeobox transcriptional factors^7,21^. It is well-established that transcriptionally active regions of the genome are enriched at the nuclear center while silent regions remain at the nuclear periphery^22^. Furthermore, LADs are mostly associated with regions of heterochromatin marked by H3K9me3^23^. As such, the physical proximity of active *HOXA* cluster to a LAD anchored by the *GLC* seems counterintuitive. However, recent studies have shown that nuclear pore complexes (NPC) embedded at the nuclear periphery can associate with both enhancer and polycomb enriched regions^24,25^. In our effort to map NPC binding on chromatin, we observed a significant enrichment of the complex at both the active distal *HOXA* region as well as the *GLC* in OCI-AML3 cells (Supplemental Figure S7c). This implies that the *GLC*-*HOXA* interaction may be the 3D chromatin organization signature of tethering active *HOXA* to nuclear pores on the nuclear periphery, promoting rapid expression of the transcription factors in HSPC and leukemias.

## Method

### RNP based CRISPR deletion of *GLC* region

RNP based CRISPR deletion was performed followed by the previous protocol ^26^. RNP complex was prepared by mixing 2μg Cas9 and 1μg sgRNA targeting each side of region of interest. RNP complex was incubated at 37 degree for 15 minutes before the electroporation. 2-3×10^5^ cells were used for one electroporation by NEON transfection system. CD34+ cells were electroporated with Buffer T under the condition “1600V, 10ms, 3 pulses”. Leukemia cell lines OCI-AML3, MV4-11 and Kasumi-1 were electroporated under the condition with Buffer R under the condition “1350V, 35ms, 1 pulse”. Deletion of targeted region was generally checked 12-16 hours after electroporation. Both purchased chemical modified guide RNA (Synthego) and in vitro transcribed guide RNA were used in this study. The target region sequence of guide RNA is listed below:

GLC-5’-sgRNA-target (chemical modified guide RNA) : AAAAGACACACCGGCGTG

GLC-3’-sgRNA-target (chemical modified guide RNA) : TCAGGAGGAAGGAGAACC

GLC-CTCF-1-sgRNA-5’ (oligos for IVT):

TtaatacgactcactataGGAAAAGACACACCGGCGTGgttttagagctagaaATAGC

GLC-CTCF-1-sgRNA-3’ (oligos for IVT):

TtaatacgactcactataGGCTGACTCGCGGGTGGGGAgttttagagctagaaATAGC

GLC-CTCF-2-sgRNA-5’(oligos for IVT):

TtaatacgactcactataGGGGGTGGAGATGGTGAGAAgttttagagctagaaATAGC

GLC-CTCF-2-sgRNA-3’(oligos for IVT):

TtaatacgactcactataGGGCGGGTTTCTCCAAGGAAgttttagagctagaaATAGC

3’HS-1-CTCF-sgRNA-5’ (oligos for IVT):

TtaatacgactcactataGGaGTCTTGGGATGGCTGAAGgttttagagctagaaATAGC

3’HS-1-CTCF-sgRNA-3’ (oligos for IVT):

TtaatacgactcactataGGtCCAAGGCAGGACATGTGTgttttagagctagaaATAGC

*HBB*-3’ enhancer-5’-sgRNA (oligos for IVT):

taatacgactcactataGGTAGGTAGATGCTAGATTCgttttagagctagaaatagc

*HBB*-3’ enhancer-3’-sgRNA (oligos for IVT):

taatacgactcactataGGAGAGCTAGTTCAAACCTTgttttagagctagaaatagc

GLC-WT-F gacagaaacttcccaggatgg

GLC-WT-R gggttggtgagattagccataaa

GLC-deletion-F (same as GLC-WT-5’) gacagaaacttcccaggatgg

GLC-deletion-R atgggatgagcaaatggaaatg

GLC-CTCF1-F: GACAGAAACTTCCCAGGATGG

GLC-CTCF1-R: GACCAGGGTTGGTGAGATTAG

GLC-CTCF2-F: AGTGCCAGTATGTTCTCACTTC

GLC-CTCF2-R: GTACGACCTCACGTGTTCAG

*HBB*-3’ enhancer -F: gaCAAGGACCACTTGAGACTC

*HBB*-3’ enhancer -R: CTGCCTATCAGAAAGTGGTGG

### HSPC *ex vivo* expansion and colony forming assay

CD34+ cord blood hematopoietic stem and progenitor cells were purchased from StemCell Tech. CD34+ cells were electroporated with 2μg Cas9 and 1μg guide RNA targeting 5’ of *GLC* and 1μg guide RNA targeting 3’ of *GLC*. CD34+ cells electroporated with 2μg Cas9 and 1μg guide RNA targeting 5’ of 3’HS-1 CTCF binding site in *HBB* locus and 1μg guide RNA targeting 3’ of 3’HS-1 were used as the control. CD34+ were recovered overnight after thawing in *ex vivo* expansion medium (SFEM II, 100ng/mL SCF, TPO, FLT3L), cells were then electroporated every 16 hours for three times. The deletion of *GLC* was checked after three electroporation. Cells from GLC and 3’HS-1 deletion was used for flow cytometry assay with CD34 (FITC, clone 581, BioLegend) and CD38 (PE, clone HB-7, BioLegend) for the check of cell differentiation. 200 GLC deleted and 3’HS-1 deleted CD34+ cells were then plated in 6 well plate with 1mL of MethoCult™ H4034 Optimum (StemCell technology). Colonies were counted 7 days after plating and the colony pictures were taken at the same time.

### CRIPSR-deletion competition assay

OCI-AML3, MV4-11 cells were electroporated with 2μg Cas9 and 1μg guide RNA targeting 5’ of *GLC* and 1μg guide RNA targeting 3’ of *GLC*. Cells were placed in the continuous culture. On day 14 the 20% of cultured cells were collected and digested with DirectPCR Lysis reagent (Viagen) and Proteinase K(Viagen), and on day 21, 20% of cultured cells were collected and digested. PCR were performed detecting the wild-type and deletion.

### Hi-C library generation for cell line and primary human AML

OCI-AML3, MV4-11 and Kasumi-1 cell lines Hi-C libraries were constructed with Arima Hi-C kit according to the manufacturer’s instruction. 1-2 million cells are used for the construction of Hi-C libraries. Each sample was sequenced to low depth to preform quality control check with Hi-C-Pro before sequencing to high depth ^27^.

Primary human AML samples were purchased from Stem Cell & Xenograft Core at University of Pennsylvania from de-identified frozen mononuclear cells from AML patients. Samples were obtained from the Stem Cell and Xenograft Core of the University of Pennsylvania. Samples were obtained after IRB consent and provided to us as annotated, anonymous samples. Mononuclear cells were thawed by directly pipetting ice cold PBS+2% BSA to frozen cells in the freezing vials under room temperature. Thawed mononuclear cells were then gradient separated with Lymphoprep (StemCell Tech). Buffy coat was then collected to get the alive cells. Cells were stained for CD33 (PE, clone WM53, BioLegend), CD34 (APC, clone 581, BioLegend), CD45 (FITC, clone HI30, BioLegend), and CD38 (PeCy7, clone HB-7, BioLegend) to check for immunophenotyping by flow cytometry. Mononuclear cells were then fixed with 2% Formaldehyde for 10mins and quenched by 0.125M glycine for 5mins. *In situ* Hi-C is performed with fixed mononuclear cells following the previous protocol ^13^. Cells were digested by DpnII (NEB) for 2 hours, end filling with biotin-dATP (Jena Bioscience) and then ligated overnight. Ligated DNA was isolated by Pheno-chloroform extraction and ethanol precipitation with phase lock tube (Qiagen). Isolated DNA was then sonicated to 200-600bp with Covaris E220 and enriched for biotin containing ligated fragments using streptavidin magnetic beads (ThermoFisher). Enriched biotin containing ligated fragments were then made to library using Accel-NGS 2S kit (Swift Bioscience).

### CRISPR tilling screening of *GLC* region

20bp oligonucleotide pool with guide RNA sequence tilling *GLC* region was synthesized by CustomArray. Oligonucleotide pool was then cloned using Gibson assembly (NEB) into BsmBI cut lentiviral vector ipUSEPR, which contains RFP and Puro as the selection marker. 293T cells were then transfected with pooled guide RNA libraries and helper plasmid pSMD2.G and pSAX2 for the production of lentivirus carrying pooled guide RNA. Unconcentrated medium is collected for the lentivirus carrying guide RNA pool. OCI-AML3 cells and MV4-11 cells were then transduced with titrated pooled guide RNA lentivirus. The titration that gave transduction efficiency between 20 and 30% was selected for large scale transduction, to ensure the M.O.I is lower than 1.

OCI-AML3 and MV4-11 cells were transduced with pLenti-Cas9-Blast lentivirus (Addgene: 52962). Transduced cells were maintained in 10μg/mL blasticidin (ThermoFisher). Single clones of the OCI-AML3-Cas9 and MV4-11-Cas9 cells were selected and maintained in 10μg/mL blasticidin. To perform large scale transduction, 10 million OCI-AML3-Cas9 and MV4-11-Cas9 cells were transduced with lentivirus carrying pooled guide RNA. 2 days after transduction, 1μg/mL puromycin (ThermoFisher) is added to cell culture. One day after puromycin selection, dead cells were removed by gradient density separation with Lymphoprep (StemCell Tech). Live cells were then kept under puromycin and blasticidin for 15 days. Cells are sampled at day 0 and day 15 of the passage to survey the *GLC* guide RNA pool. Genomic DNA was extracted from cells in culture at day 0 and day15. Guide RNA pool was amplified with the following primer F-CTTGTGGAAAGGACGAAACACCG, R-CCTAGGAACAGCGGTTTAAAAAAGC. Illumina sequencing indexes were added by PCR amplification and indexed libraries were sequenced on HiSeq2000 by Fulgent. Guide RNA with specificity score less than 35 was filtered out, and the remaining guide RNA sequence was aligned to *GLC* genomic location. The abundance of certain guide RNA was normalized in day 0 and day 15 library. The drop-off and enrichment score was calculated as *log*10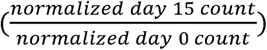

### ChIP-seq

ChIP in OCI-AML3 and MV4-11 cell lines are performed with previously described ChIPmentation protocol ^28^. One million cells were fixed with 1% formaldehyde and sonicated with Covaris E220. Sonicated chromatin were then incubated with antibody against H3K27me3(Cell Signaling, #9733), Nuclear Pore Complex (Abcam, Mab414, ab24609) and CTCF (Abcam, #ab70303) overnight. Protein A/G beads (ThermoFisher) were added to the sonicated chromatin for 2 hours. After washes with low-salt, high-salt and LiCl wash buffer. Following washes, beads were tagmented by Illumina tagmentation enzyme mix for 10 mins at 37 °C. After the tagmentation reaction stopped and the wash of beads, pull-downed DNA was de-crosslinked with Proteinase K (Zymo Research) at 55 °C for 3 hours. Pull-downed DNA was extracted by DNA clean and concentration kit (Zymo Research). PCR amplification was performed, and libraries were sequenced with Illumia NextSeq Platform.

### FISH imaging

FISH probes were purchased from Empire Genomics. The BAC clone RP11-1025G19 was used for the labelling of *HOXA* locus (*HOXA-AS3* to *EVX1*). The BAC clone RP11-598H18 was used for the labelling of *GLC* locus. FISH was performed with the modification of protocol used previously ^29^. Cells were fixed with 4% paraformaldehyde (Sigma-Aldrich) and spread on Superfrost Plus microslides (VWR). Coverslips were incubated in 20% glycerol in PBS for one hour at room temperate, dipped in liquid nitrogen for 6 time, washed three times with PBST in coplin jar for 5 minutes each, rinsed and incubated on ice with 0.1 M HCl/0.7% Triton X-100/2× saline-sodium citrate (SSC) for 10 min. The coverslips were then washed with 2× SSC for 3 times in coplin jar for 5 minutes each. 2μL FISH probes were then added to coverslips. Coverslips are then mounted and sealed with rubber cement. Samples were denatured on Visys HYBRITE heat block at 76°C for 3 min and then hybridized 18–72 h on Visys HYBRITE heat block at 37°C.

Coverslips were washed at 37°C for 5 min three times in 2× SSC and then at 60°C for 5 min three times in 0.1× SSC and then at room temperature in 4× SSC for 5 min. Coverslips were mounted in DAPI-containing antifade mounting media. The image was taken by Nikon A1plus-RSI confocal microscope.

### Immuno-FISH imaging

For anti-LaminB Immuno-FISH, cells were firstly fixed in 3% paraformaldehyde in PBS for 10 min at room temperature and washed in PBST for 5min 3 times. Cells were then permeabilized with 0.1% Triton-X at room temperature for 10 mins, then washed in PBST for 5min 3 times. Cells were then incubated with rabbit anti-LaminB antibodies diluted 1:1,000 in blocking buffer overnight at 4°C, washed with PBST 5 min for three times at room temperature, incubated for 1 hour at RT with Alexa488-labeled secondary goat anti-rabbit IgG (ThermoFisher.). Cells were then proceeded to FISH protocol mentioned above for washes and pretreatment. After 2XSSC washes, Coverslips were incubated at 2xSSC overnight at 37 °C before the FISH probes were added. Coverslips were then proceeded to FISH protocol mentioned above for the FISH hybridization.

### Loop calling

Hi-C Computational Unbiased Peak Search is used to call loops from Hi-C contact maps of CD34+ and GLC-deleted cell lines. The loops are called at 25 kb resolution. Other parameters are set as default values, namely false discovery rate (FDR) = 0.1, window size = 3 and peak-merging threshold = 50 kb.

HICCUPS compares the contact reads of each pixel with its four nearby regions named horizontal, vertical, donut and lower-left, which correspond to the horizontal / vertical lines crossing at the pixel, the doughnut-shaped surrounding region and the lower-left quarter of the ring, respectively. Then a Benjamini-Hochberg FDR control procedure is applied to filter the significantly enriched pixels. Only if the four regions’ q-values are all below the FDR threshold can a region be identified as a loop.

## Data access

All processed data for visualization can be accessed at the following URL https://www.dropbox.com/sh/6wu95419kfwaenj/AADMDgwAB4RK6GqIFH0meeUZa?dl=0

### Public data used in this study

GEO accession number GSM3024909 for OCI-AML3 H3K27ac ChIP-seq; GEO accession number GSM1534446 for Kasumi-1 H3K27me3 ChIP-seq; GEO accession number GSM2212053 for Kasumi-1 H3K27ac ChIP-seq; GEO accession number GSM1525022 for OCI-AML3 whole genome bisulfite sequencing; GEO accession number GSM486704 for CD34+ HSPC H3K27me3; GEO accession number GSM916052 for CD34+ HSPC whole genome bisulfite sequencing; GEO accession number GSM772870 for CD34+ HSPC H3K27ac.

## Author Contribution

X.Q.D.W and X.Z conceptualized the idea and performed experiments, analyzed the data. W.Z and P.H analyzed Hi-C data. H.G generated Hi-C and ChIP-seq libraries. Y.L performed FISH and immuno-FISH staining. Y.L and C.C designed guideRNA library targeting *GLC.* F.F and J.L analyzed and quantified the 3D genomic interaction data. G.P provide advice and jointly supervise the project with X.Z. All authors participated in the writing and editing of the manuscript.

## Acknowledgement

We acknowledge Drs. Margaret Goodell and Mira Jeong for the initial contribution to the project. We thank for Drs. Yun Huang, Stanley Lee and Yali Dou for the help of reviewing the manuscript. X.Z is supported by VARI fellowship, ASH scholar award and is an EvansMDS Young Investigator.

