## Supplementary material for "Three-dimensional regulation of *HOXA* cluster genes by a *cis*-element in hematopoietic stem cell and leukemia": Supplemt Figure 1-7

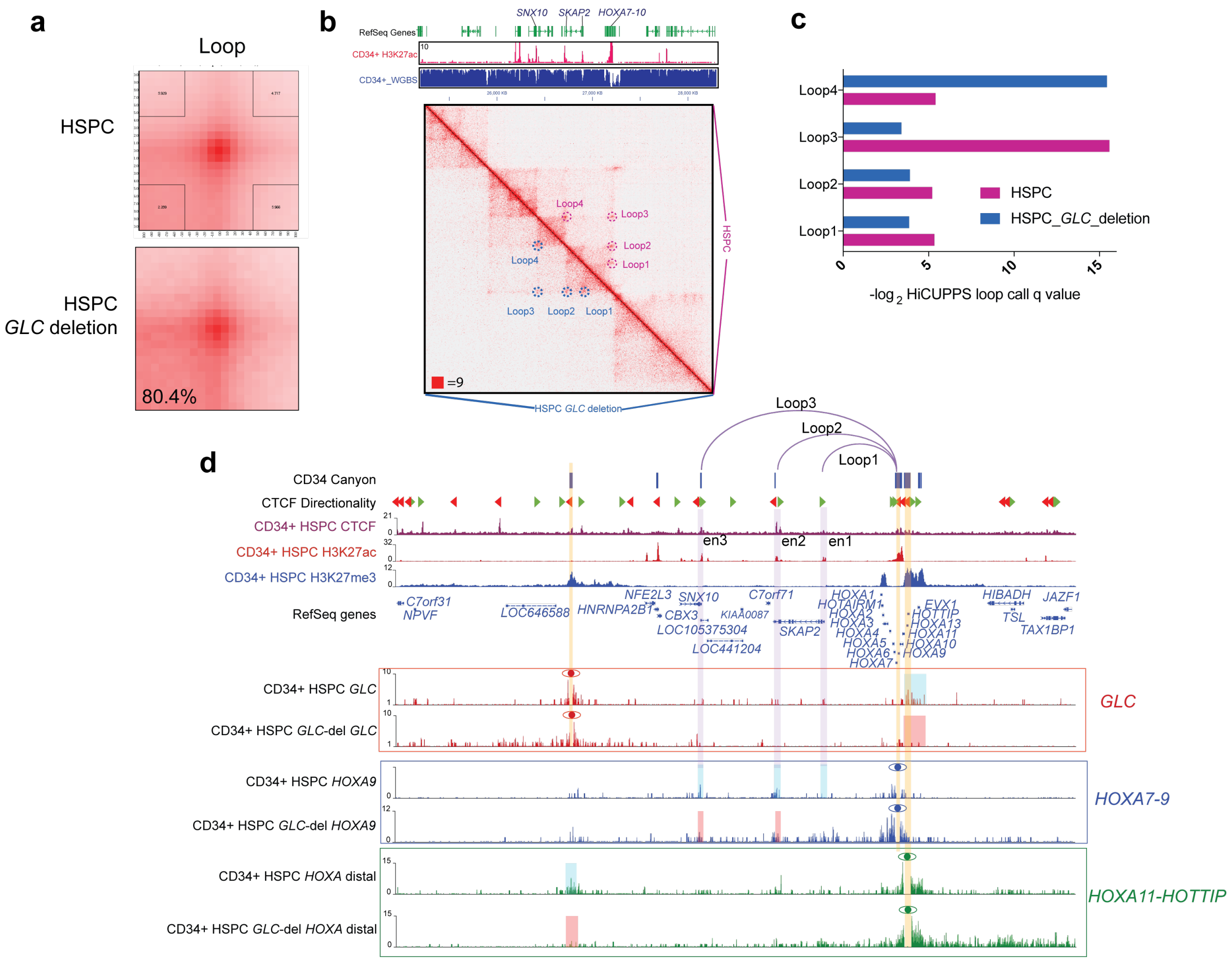

**Supplemental Figure 1. The loop quantification in the HSPC with partial *GLC* deletion.**

**a).** The APA analysis of CD34+ HSPC and HSPC with *GLC* deletion. Mild global reduction of loop interaction is observed. **b).** The juicebox view of HiC contact matrix in HSPC and HSPC with *GLC* deletion showing the loop interaction called in either samples. Loop location is annotated. **c).** The HiCUPPS loop calling q value of loops annotated in **(b)** of 2 samples. A value indicates the strength of loop signal over the background. **d).** The virtual 4C view of the interaction from H3K27me3 covered *GLC*, H3K27me3 covered *HOXA11-HOTTIP* and H3K27ac covered *HOXA7-9* region in CD34+ WT and CD34+ *GLC*-deletion HSPC. Yellow bar indicates the *GLC* region and *HOXA7-9* region used as viewpoint in virtual 4C. Blue bars show the regions interacts with the viewpoint in CD34+ WT and red bars show the regions that interact with viewpoint region in CD34+ HSPC after *GLC* CRISPR deletion. Purple bar highlights *SNX10* (en3), *SKAP2* TSS (en1), *SKAP2* TES (en2) enhancer region. Purple arc highlights the interaction region between *HOXA7-9* viewpoint and *SNX10*, *SKAP2* TSS, *SKAP2* TES enhancer through convergent CTCF interactions.

**a**

| Collection_ID | Subtype | WBC | Blasts | Blast % | Gene | MutationType | Protein_Change | VAF | Blast flow |
|---|---|---|---|---|---|---|---|---|---|
| 4943 | AML with 11q23 abnormalities | 109.2 | 91 | 83.33333 | PHF6 | deletion | p.M46Ifs*35 | 96.12 | CD5- CD10- CD13- CD14- CD15+ CD19- CD20- CD33+ CD34- CD56 dim + CD64+ CD79- HLA-DR+ TdT- |
| 5577 | M1 [without maturation] | 142.8 | 82 | 57.42297 | FLT3 | insertion | p.Y599_D600insGMTGSSDNEYFYVDFR | 25.87 | CD5- CD10- CD13 dim + CD14- CD15- CD19- CD20- CD33+ CD34- CD38 var + CD56- CD64 minor subset + CD79- HLA-DR- TdT- |
| | | | | | IDH1 | substitution | p.R132G | 48.76 | |
| | | | | | NPM1 | insertion | p.W288Cfs*12 | 46.81 | |
| 6527 | AML-MLD hx prior MDS/MPN | 170.7 | 75 | 43.93673 | CBL | substitution | p.C404Y | 51.77 | CD5- CD10- CD14- CD19- CD20- CD33+ CD34- CD56+ |
| | | | | | FLT3 | insertion | p.F612_G613insSDFRDYEYDLKWEFPR | 18.16 | |
| | | | | | NPM1 | duplication | p.W288Cfs*12 | 33.56 | |
| | | | | | TET2 | deletion | p.S1448Vfs*10 | 46.49 | |
| | | | | | TET2 | deletion | p.T710Mfs*41 | 44.99 | |
| | | | | | CBL | substitution | p.C404Y | 49.42 | |
| | | | | | FLT3 | insertion | p.F612_G613insSDFRDYEYDLKWEFPR | 21.9 | |
| | | | | | NPM1 | duplication | p.W288Cfs*12 | 37.02 | |
| | | | | | TET2 | deletion | p.T710Mfs*41 | 46.33 | |
| | | | | | TET2 | deletion | p.S1448Vfs*10 | 48.62 | |

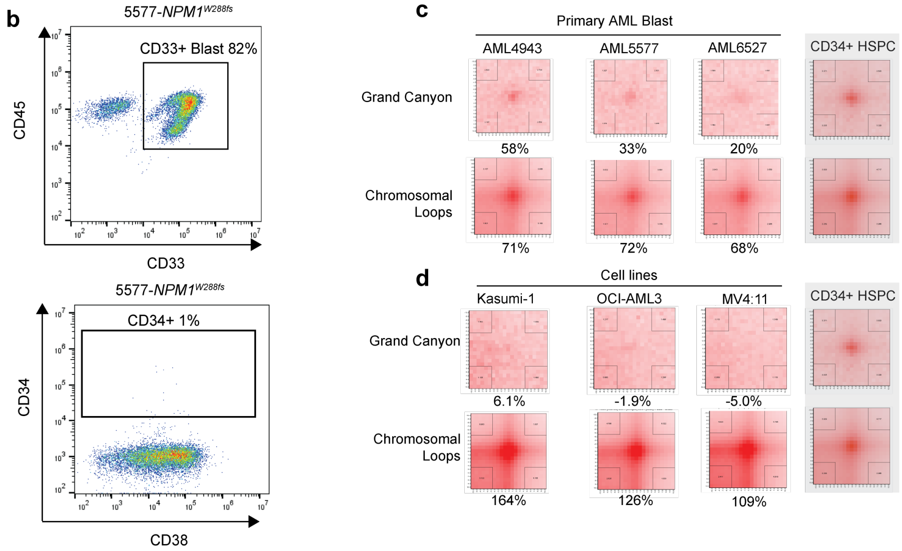

**Supplemental Figure 2. The characteristic of primary AML blast and the global APA analysis.**

**a**). The mutation landscape of AML patient blasts used in this study. **b**). The flow cytometry plot of immunophenotype of AML blast from AML-5577. CD33, CD34 and CD38 is used to determine the immunophenotype of AML-5577 blast. **c**). The aggregation peak analysis (APA) for the CD34+ CD38- grand canyon and chromosomal loop interaction in primary AML blasts used in this study. **d**). The aggregation peak analysis (APA) for the CD34+ CD38- grand canyon and chromosomal loop interaction in AML cell lines used in this study.

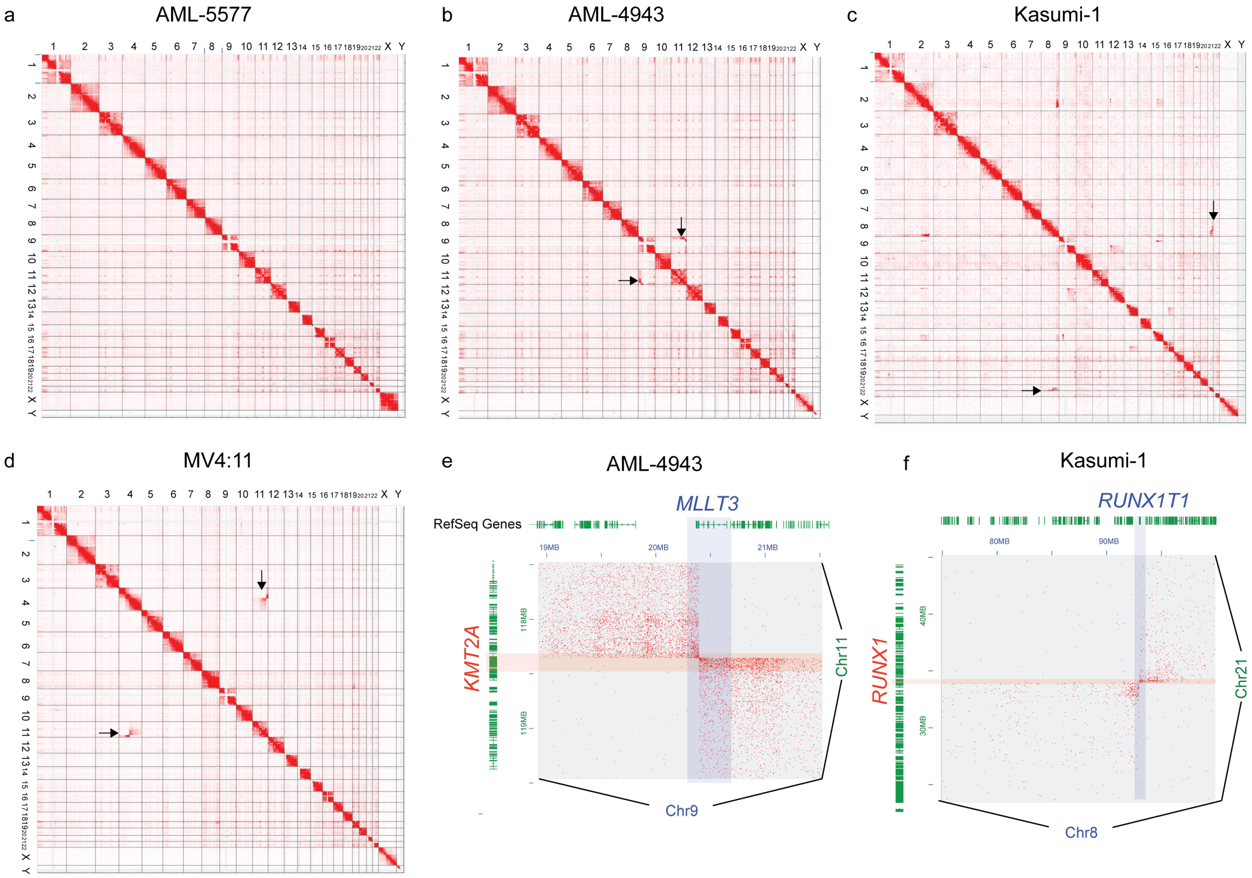

**Supplemental Figure 3. Translocation in primary and AML cell line detected by HiC.**

**a)**. The interaction matrix of all chromosomes from cytogenic normal AML-5577. **b)**. The interaction matrix of all chromosomes from cytogenic normal AML-4943. The arrows indicate the translocation signal between chromosome 9 and chromosome 11. **c)**. The interaction matrix of all chromosomes from Kasumi-1 cell line, which is known for carrying t(8:21) translocation. The arrows indicate the translocation signal between chromosome 8 and chromosome 21. **d)**. The interaction matrix of all chromosomes from cytogenic normal MV4:11 cell line, a cell line carrying t(4:11) translocation. The arrows indicate the translocation signal between chromosome 4 and chromosome 11. **e).** The kilobase level translocation signal map of t(9:11) translocation in AML-4943. Red shade highlights the genomic location of *KMT2A* gene on chromosome 11. Blue shade highlights the genomic location of *MLLT3* gene on chromosome 9. **f).** The kilobase level translocation signal map of t(8:21) translocation in Kasumi-1. Red shade highlights the genomic location of *RUNX1* gene on chromosome 21. Blue shade highlights the genomic location of *RUNX1T1* gene on chromosome 8.

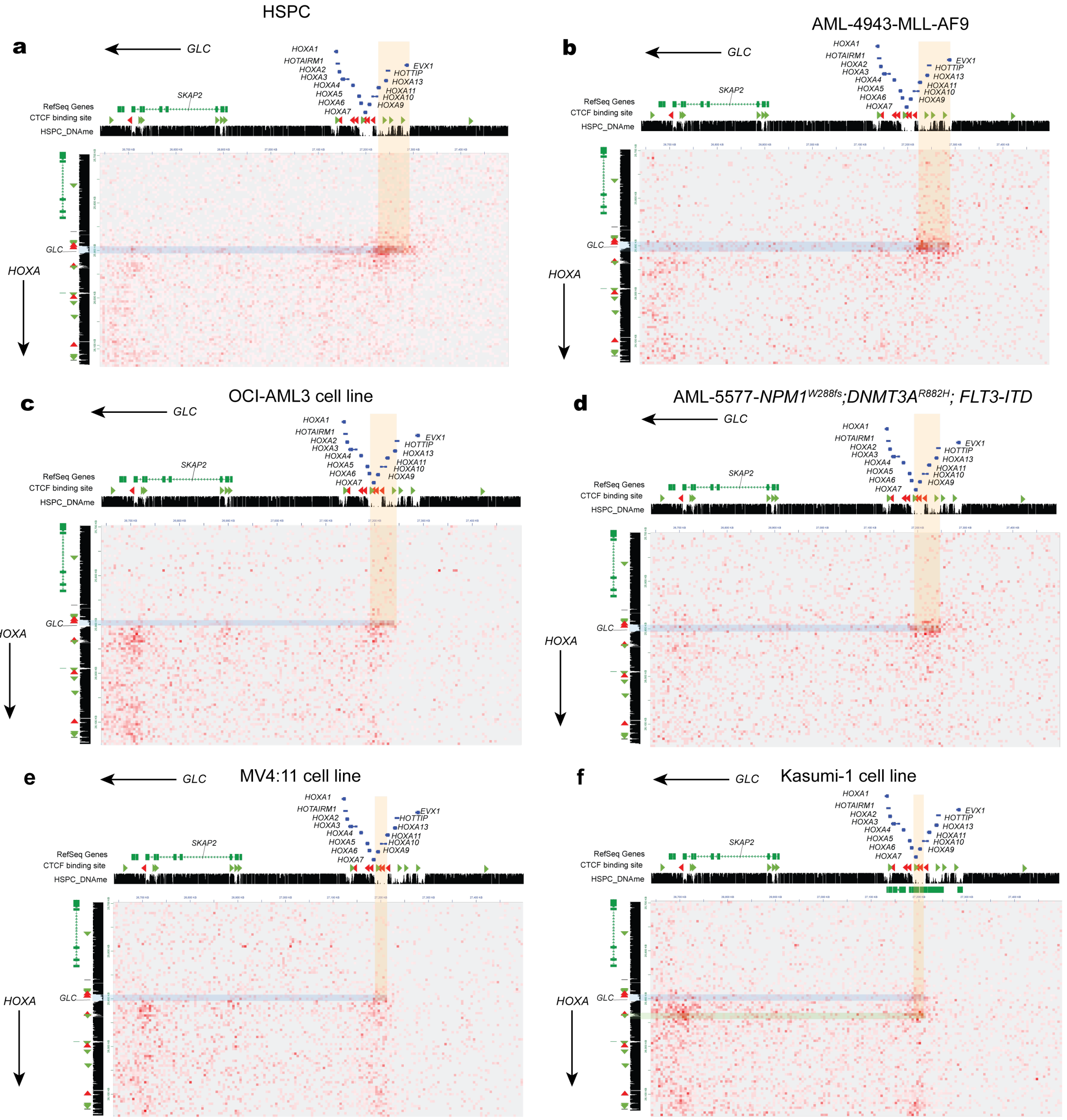

**Supplemental Figure 4. *GLC-HOXA* cluster interaction in AML and cell lines.**

**a).**The zoomed-in juicebox view of the interaction between *GLC* and *HOXA* cluster in CD34+ CD38- HSPC. Green arrows represent the CTCF binding site with reverse orientation and red arrows represent the CTCF binding site with forward orientation. Blue shade highlights the *GLC* region and yellow region highlights the *HOXA11* to *HOTTIP* and *EVX1* region. Note the CTCF sites in *HOXA* region associated with the interaction of *GLC* are not in convergent orientation **b).** The zoomed-in juicebox view of the interaction between *GLC* and *HOXA* cluster in AML-4943 Blast. Blue shade highlights the *GLC* region and yellow region highlights the *HOXA11* to *HOTTIP* and *EVX1* region. **c).** The zoomed-in juicebox view of the interaction between *GLC* and *HOXA* cluster in OCI-AML3 cell lines. Blue shade highlights the *GLC* region and yellow region highlights the *HOXA7* to *HOTTIP* region. **d).** The zoomed-in juicebox view of the interaction between *GLC* and *HOXA* cluster in AML 5577 blast. Blue shade highlights the *GLC* region and yellow region highlights the *HOXA7* to *HOXA11* region. **e).** The zoomed-in juicebox view of the interaction between *GLC* and *HOXA* cluster in MV4:11 cell line. Blue shade highlights the *GLC* region and yellow region highlights the *HOXA9* to *HOXA11* region. **f).** The zoomed-in juicebox view of the interaction between *GLC* and *HOXA* cluster in MV4:11. Blue shade highlights the *GLC* region and yellow region highlights the *HOXA7* to *HOXA9* region. Green shade highlights the down stream 5' CTCF sites to *GLC*. The CTCF sites forms CTCF convergent cohesion/CTCF loop with *SKAP2* transcription end site and *HOXA7-HOXA9* region.

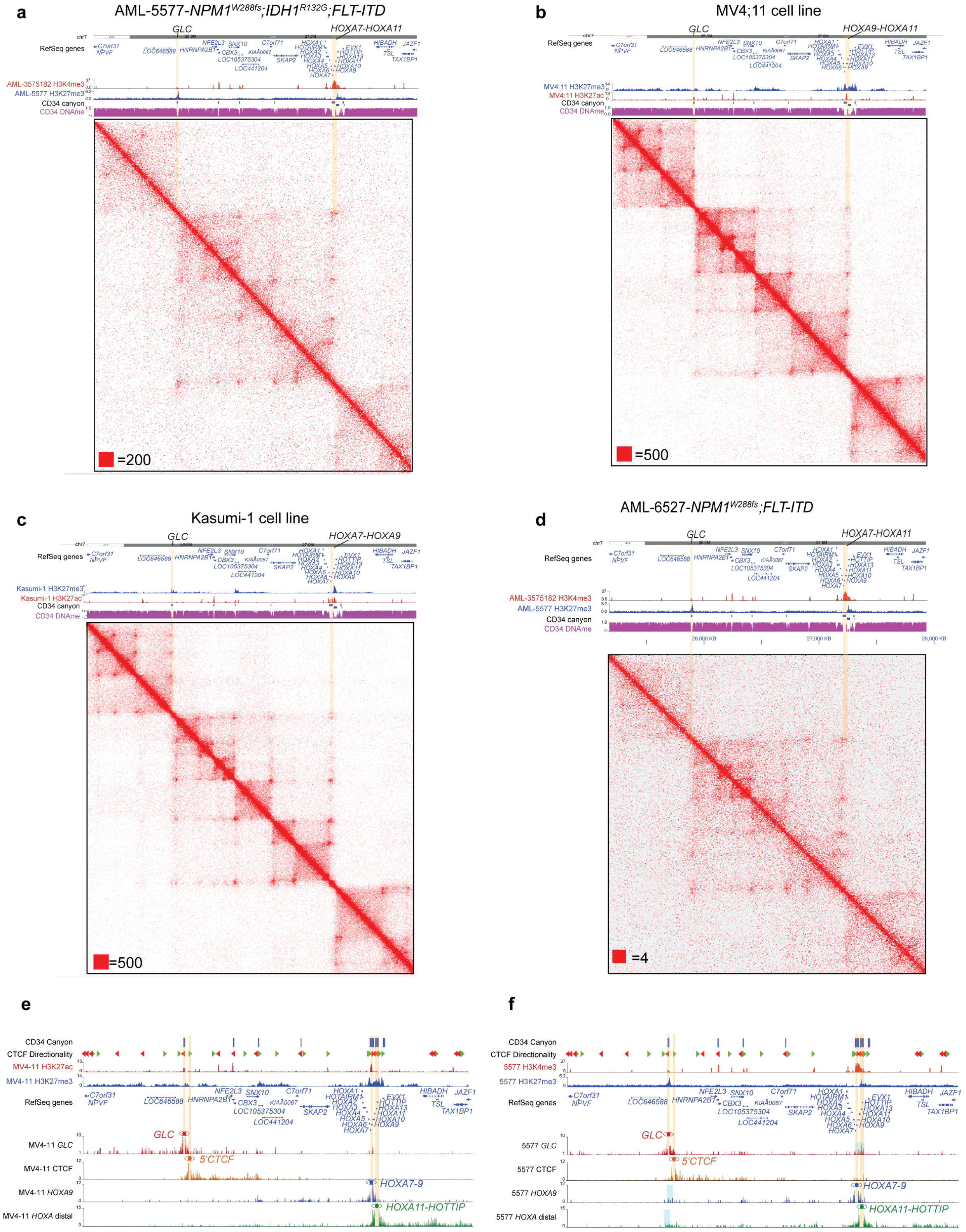

**Supplemental Figure 5. HiC interaction map of primary AML blast and AML cell lines.**

**a**). The contact heatmap in 3 TAD regions in around *HOXA* cluster in primary leukemia blast AML5577. AML5577 carries the mutation in *NPM1, IDH1* and *FLT3*. Yellow shade indicates the interacting regions of *GLC* and corresponding *HOXA* regions (*HOXA7-HOXA11*). **b**). The contact heatmap in 3 TAD regions in around *HOXA* cluster in MV4:11 cell lines. Yellow shade indicates the interacting regions of *GLC* and corresponding *HOXA* regions (*HOXA9* to *HOXA11*). **c**). The contact heatmap in 3 TAD regions in around *HOXA* cluster in Kasumi-1 cell lines. Yellow shade indicates the interacting regions of *GLC* and corresponding *HOXA* regions (*HOXA7* to *HOXA9*). **d**). The contact heatmap in 3 TAD regions in around *HOXA* cluster in AML-6527 AML blasts. AML-6527 carries mutations in *NPM1, TET2* and *FLT3.* Yellow shades indicate the interacting regions of *GLC* and corresponding *HOXA* regions (*HOXA7* to *HOXA11*). **e).** The virtual 4C interaction profile from the viewpoint of *GLC*, 5' CTCF, *HOXA7-9, HOXA11-HOTTIP* site of MV4:11 cell line. Yellow strips highlight the viewpoint used region for virtual 4C. **f).** The virtual 4C interaction profile from the viewpoint of *GLC*, 5' CTCF, *HOXA7-9, HOXA11-HOTTIP* site of AML-5577-Blast. Yellow strips highlight the viewpoint used region for virtual 4C. The blue bars highlight the regions that interacts with viewpoint region.

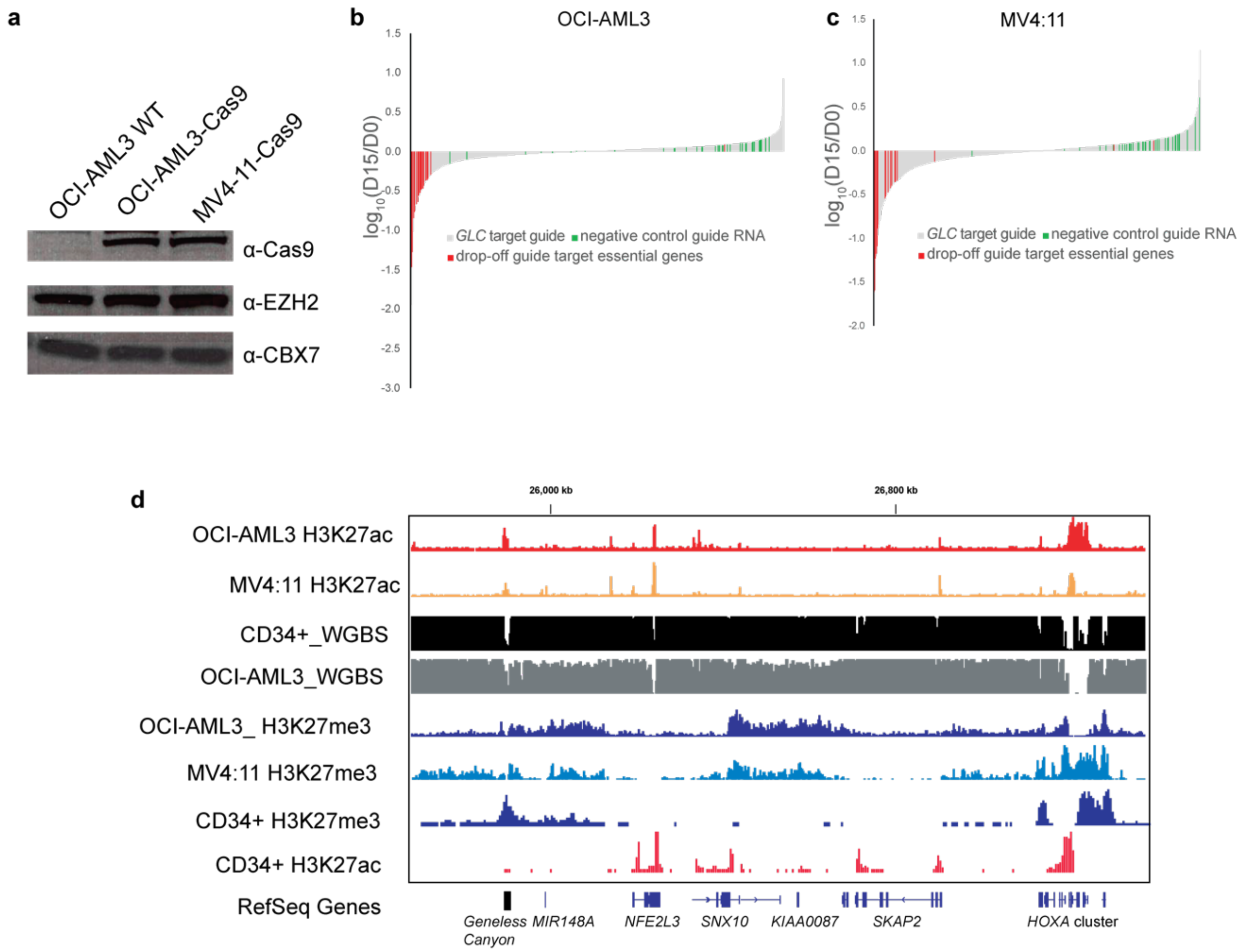

**Supplemental Figure 6. The *Geneless Canyon* CRISPR/Cas9 tilling screen.**

**a).**The stable expression of Cas9 in OCI-AML3 cell line and MV4:11 cell. **b-c).** The waterfall plot of guide RNA abundance change in the *GLC* tilling CRISPR/Cas9 mutagenesis screen in OCI-AML3 cells **(b)** and MV4:11 cells **(c)**. Red bars represent the guide RNAs that target the essential genes (*POL2A, RPS20, RPL9, RPL23A, RPA3, PCNA, POLR2A, POLR2D, PCNA* ) and genes driving cell growth (*MYC, CDK1, BRD4*) in the *Geneless Canyon* screen library pool. Green bars represent the negative control guides (*Luc, Ren, LacZ, ROSA26*) that do not target human genome. **d).** The epigenomic landscape between *Geneless Canyon* and *HOXA* cluster in the OCI-AML3 and MV4:11 cells. The *Geneless Canyon* region is now activated with H3K27ac mark in both OCI-AML3 and MV4:11 cells.

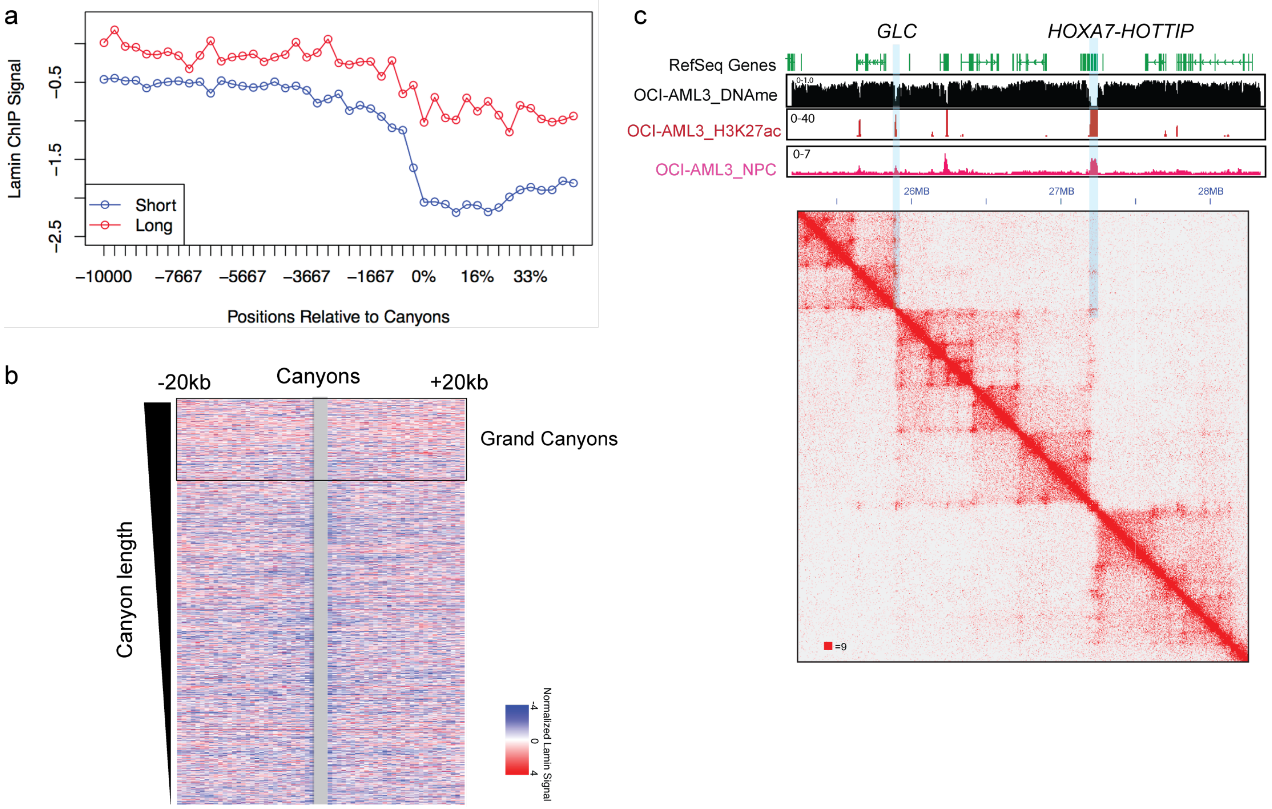

**Supplemental Figure 7. The spatial proximity of grand DNA methylation canyons to LAD and nuclear pores.**

**a).** Lamin B ChIP signal distribution in canyon regions[1]. **b).** Lamin B ChIP signal in the regions surrounding DNA methylation canyons. The normalized DNA methylation canyon regions are highlighted in grey. Grand canyons are defined as DNA methylation canyon with length over 7.3kb. **c).** The juicebox view of 3D genomic interaction in OCI-AML3 cells in the 3 TADs surrounding *HOXA* cluster. The H3K27ac [2] and OCI-AML3 nuclear pore complex (NPC) ChIP-seq track is overlaid on top. The blue stripe highlighted the *GLC* and *HOXA7-HOTTIP* region, which in OCI-AML3 cell lines are strongly enriched for H3K27ac and NPC binding.
